# Evaluating predictive biomarkers for a binary outcome with linear versus logistic regression – Practical recommendations for the choice of the model

**DOI:** 10.1101/347096

**Authors:** Damian Gola, Nicole Heßler, Markus Schwaninger, Andreas Ziegler, Inke R. König

## Abstract

A predictive biomarker can forecast whether a patient benefits from a specific treatment under study. To establish predictiveness of a biomarker, a statistical interaction between the biomarker status and the treatment group concerning the clinical outcome needs to be shown. In clinical trials looking at a binary outcome, linear or logistic regression models may be used to evaluate the interaction, but the effects in the two models are different and differently interpreted. Specifically, the effects are estimated as absolute risk reductions (ARRs) and odds ratios (ORs) in the linear and logistic model, thus measuring the effect on an additive and multiplicative scale, respectively.

We derived the relationship between the effects of the linear and the logistic regression model allowing for translations between the effect estimates between both models. In addition, we performed a comprehensive simulation study to compare the power of the two models under a variety of scenarios in different study designs. In general, the differences in power to detect interaction were minor, and visible differences were detected in rather unrealistic scenarios of effect size combinations and were usually in favor of the logistic model.

Based on our results and theoretical considerations, we recommend to 1) estimate logistic regression models because of their statistical properties, 2) test for interaction effects and 3) calculate and report both ARRs and ORs from these using the formulae provided.

## Introduction

Novel technologies and increased accumulated knowledge on the functional background of diseases have made the application of biomarkers in clinical studies increasingly popular. Their use is extremely diverse and includes serving as a tool for diagnosis, for staging the disease, for forecasting disease prognosis or for monitoring and predicting clinical response [1]. For many instances, it is most helpful to distinguish between prognostic biomarkers and predictive biomarkers [2].

Prognostic biomarkers can forecast the development of the disease. In a randomized clinical trial, this would usually be the outcome of the study such as remission. Importantly, this forecast is independent of the intervention but an overall prognosis. Put differently, patients with different prognostic biomarker profiles would have a different course of disease, regardless of the intervention group. For example, the epidermal growth factor receptor tyrosine kinase status is a prognostic factor for survival in patients with non-small cell lung cancer [3], irrespective of the treatment. Predictive biomarkers, in contrast, predict the effect of the intervention itself and therefore serve as companion diagnostic tests [4]. Thus, patients with different predictive biomarker values would differ in how likely they are to benefit from the specific therapy under study or to suffer from side effects. For instance, several studies have shown that eosinophil counts in peripheral blood are predictors for treatment response to Anti-IL-5 in patients with severe asthma [5–7].

Biomarkers are considered in clinical trials using different study designs, and these are described in detail in the literature [2, 4, 8]. Which design should be used depends, among other aspects, mostly on what is already known about the biomarker and the overall aim of the study. If the aim is to prove the predictiveness of a biomarker, all patients regardless of their biomarker status need to be randomized to the treatment groups. This is integrated in the so-called “randomize-all” or “biomarker-stratified” design. Specifically, in the “randomize-all” design, eligible patients are randomized into the treatment groups before their biomarker status is assessed (Fig 1A). In the “biomarker-stratified randomization” design, the biomarker status is assessed first. Then, patients with positive and negative biomarker status are randomized separately (Fig 1B).

**Fig 1.**
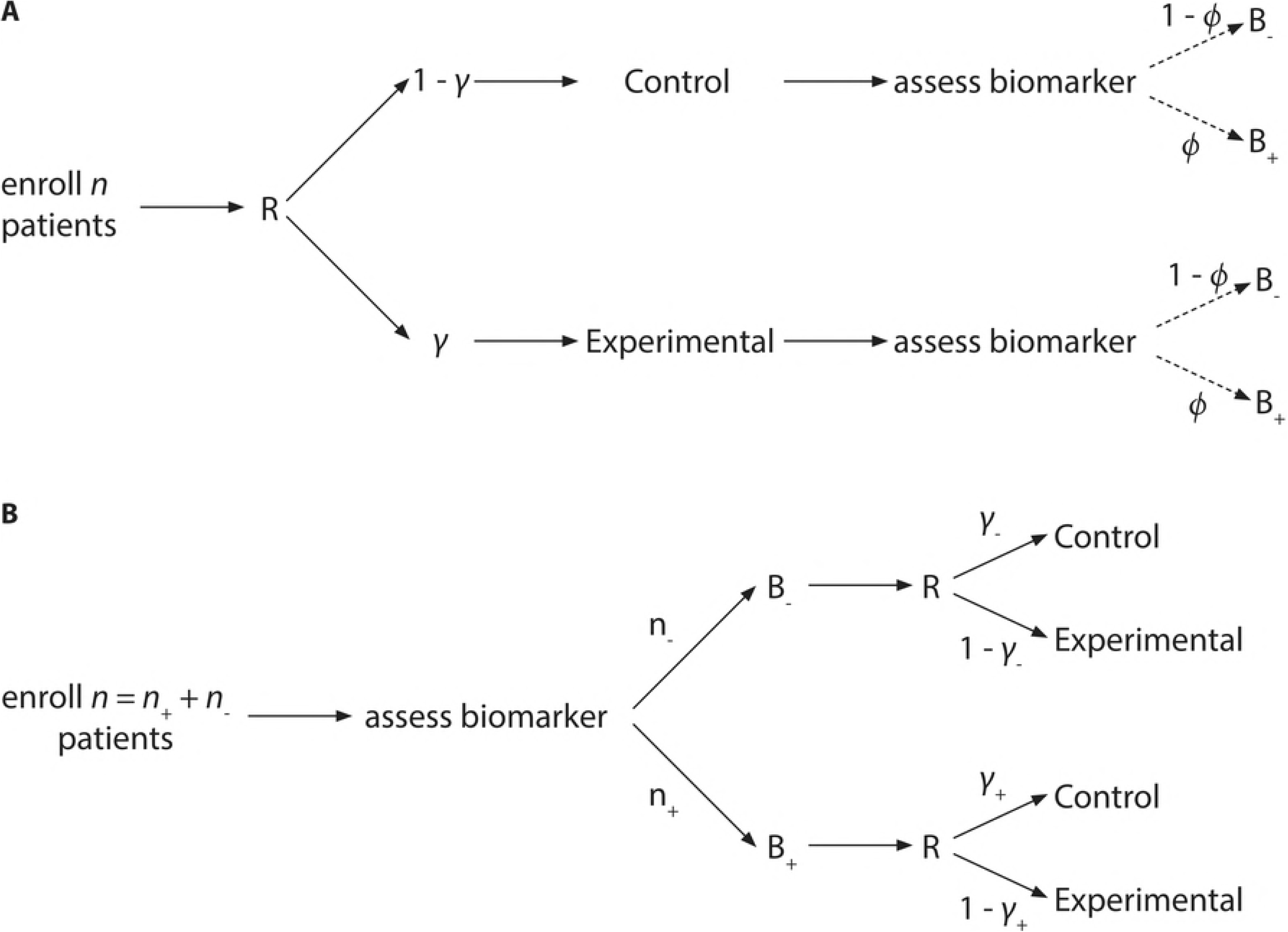
Trial designs used in the simulation study. (A) In the “randomize-all” design *n* patients are assigned irrespectively of their biomarker status to one the treatment groups based on the randomization factor *γ*. (B) In the “biomarker-stratified randomization” design, *n* patients are assigned to two randomizations based on their biomarker status.

If, in contrast, only patients with a positive biomarker status are randomized as in the “targeted” design, it can only be shown that there is a treatment effect in this group, which does not rule out that also biomarker negative patients benefit from the intervention, who were not investigated. Furthermore, for establishing a predictive biomarker the trial needs to show statistically that the treatment effect depends on the biomarker status, i.e., the interaction between treatment arm and biomarker status has to be established. However, it does not suffice to analyze biomarker positive and negative subgroups in separate trials and report an effect in one but not the other group [9]: Firstly, not finding the therapeutic effect in one group might be due to a lack of power. For example, in the study by Pant et al. [10] predictiveness of albumin for the treatment of advanced pancreatic cancer with bevacizumab was claimed on the finding of a positive effect in patients with normal baseline albumin but not in others. However, only 26 patients with non-normal serum albumin levels were included in the study. Hence, the confidence interval of the effect is very wide in this subgroup and indeed includes the effect observed in patients with normal serum albumin. Consequently, it cannot be ruled out that the effect was only not detected in the smaller group, and no interaction between the treatment and albumin can be observed. A second reason against claiming predictiveness based on the analysis of subgroups only is that even if there are effects in both subgroups, predictiveness of the biomarker cannot be excluded, because the therapeutic effect might be weaker (quantitative interaction) or in the opposite direction (qualitative interaction) in the second subgroup.

In the following, we will describe the statistical methods to evaluate the biomarker-by-treatment interaction that needs to be shown for the predictiveness of a biomarker.

## Statistical evaluation of biomarker-by-treatment interaction

The statistical method of choice to evaluate the biomarker-by-treatment interaction depends on the data, i.e., the scale of the outcome variable and additional covariables that are to be included in the model. In the following, we will focus on the simple setting of a dichotomous outcome without further covariables. As a first approach, a linear regression framework can be used in which the risk or probability of the dichotomous outcome *y* (e.g. therapy success) is modeled as a function of the dichotomous variables treatment *T*, biomarker status *B*, and treatment-by-biomarker interaction *TB* with

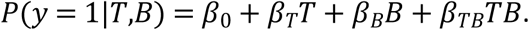

Here, *T* = 0 or *T* = 1 if a patient receives the control treatment or the experimental treatment, *B* = 0 or *B* = 1 if a patient is biomarker negative or positive, and *TB* = 1 only if a biomarker positive patient receives the experimental treatment. Through this, the coefficients *β_T_* and *β_B_* can be interpreted as the increase in risk with a change in the treatment group and the biomarker status, respectively. The interpretation of these effect estimates as absolute risk reductions (ARRs) is beneficial since it can be directly related to the number needed to treat (NNT=1/ARR) [11]. The coefficient *β_TB_* indicates whether the influence of *T* and *B* on *y* is independent, in which case it would equal 0. If it deviates from 0, there is a statistical interaction between *T* and *B* regarding the risk of the outcome on the additive scale [12].

However, this model has some statistical disadvantages. For example, the predicted probability might be out of the range of possible values between 0 and 1. The standard statistical model for analyzing dichotomous outcome in the life sciences therefore is the logistic regression model. Here, the log odds of the outcome *y* is modeled as a function of *T* and *B* and their interaction *TB* by

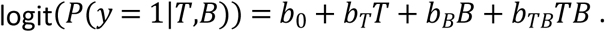

From this, the coefficients and *b_T_* and *b_B_* can be exponentiated to be interpreted as the increase in odds of the outcome with a change in the treatment group and the biomarker status, respectively. The coefficient *b_TB_*, when exponentiated, then measures the treatment-by-biomarker interaction as the odds ratio (OR) on the multiplicative scale. One advantage of this model is that the predicted outcome probability will be guaranteed to lie between 0 and 1. Furthermore, the logit link is the natural parameter from the linear exponential family which provides excellent statistical properties.

The linear and the logistic models are different, they have different effect sizes. This can be seen from S1 Appendix in which we have derived the relation between ARRs from the linear probability model and ORs from the logistic regression model.

Concerning the interaction effect, it can be shown that the models lead to different results, meaning that the evidence for interaction will differ in strength, and that interaction in one model does not imply interaction in the other. For example, in the study by Bokemeyer et al. [13], patients with metastatic colorectal cancer had been randomized to receive FOLFOX-4 with or without cetuximab and were screened for *K-ras* mutations. A randomize-all design was used, and, amongst other criteria, the best overall response in both *K-ras* positive and negative patients was analyzed separately. We re-analyzed the data presented in the paper and derived that the relative risk of response from a linear regression model under cetuximab plus FOLFOX-4 versus FOLFOX-4 only was 1.68 in the wild type and 0.64 in the mutation group, respectively. The corresponding p-value for the interaction was 0.00019. In the logistic regression model, the odds ratio of response was 2.60 in the wild type and 0.46 in the mutation group, respectively, with an interaction p-value of 0.00023. Therefore, even though interaction was established in both models, the p-values differ [13].

Therefore, given the statistical advantage of the logistic regression model over the linear probability model, one may question the use of the linear regression model in this setting in general. However, it has been shown that the statistical problems may not be as large as anticipated [12, 14] and that, considering the interpretation of the effects, there are indeed some merits to the linear model. As notional example, we consider the data in Table 1 (left), showing the risk or probability of an outcome depending on the treatment and biomarker status. In this example, changing the biomarker status from negative to positive always increases the risk by 20%, and changing the treatment from control to experimental always increases the risk by 40%. Thus, there is no additive biomarker by treatment interaction. We now assume that we wish to select patients who will benefit most from the treatment. If there were 100 patients each who were biomarker positive and negative, 10 and 30 would reach a positive outcome, respectively, under control treatment (Fig 2A). Switching to the experimental treatment instead, the numbers could be increased to 50 and 70, respectively. This means that in either biomarker group, 20 patients would benefit from the experimental treatment, indicating that the biomarker status does not need to be taken into account when offering the treatment, which is mirrored by the lack of an additive interaction. Consider now the data in Table 1 (right), where changing the biomarker status from negative to positive increases the risk by 10% under control but by 30% under the experimental therapy, and changing the treatment from control to experimental increases the risk by 20% for biomarker negative and by 40% for biomarker positive patients. Phrased differently, changing the biomarker status is always associated with doubling the risk, and changing the therapy regimen with a 3-fold increase. In this case, there is therefore no multiplicative interaction. Translating these risks into patient numbers who will benefit from the treatment (Fig 2B) now shows that by switching the treatment from control to experimental would benefit 20 biomarker-negative but 40 biomarker-positive patients. Given limited resources, it might therefore be reasonable to offer the experimental treatment preferably to biomarker positive patients, even though there is no biomarker by treatment interaction on the multiplicative scale. From a health economic point of view, it can therefore be argued that interaction on the additive scale, thus use of the linear regression model, should at least complement the logistic regression model.

**Fig 2.**
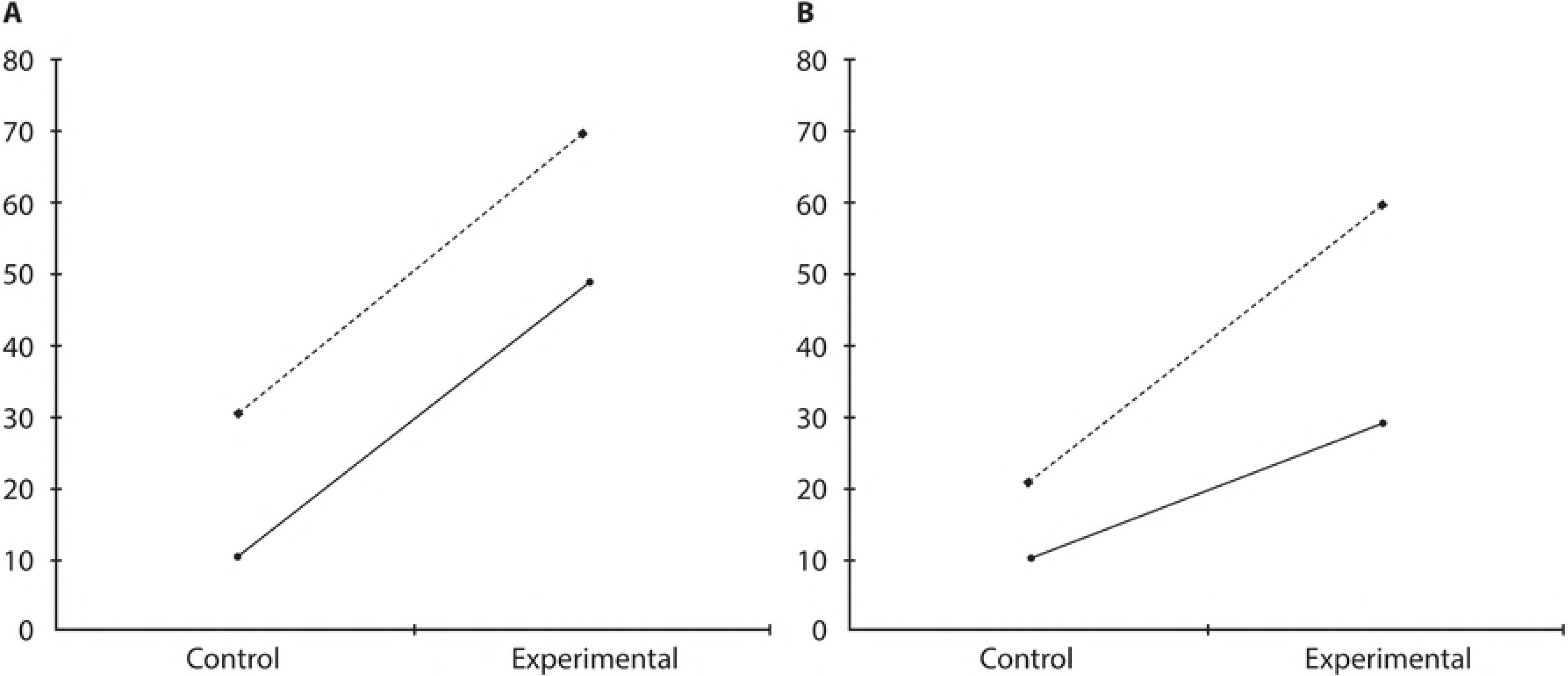
Number of patients with a positive outcome. Based on a sample size of 100 in every constellation in the scenario of (A) no additive interaction and (B) no multiplicative interaction as specified in Table 1. Solid line: biomarker negative, dashed line: biomarker positive.

**Table 1.** Notional risk of outcome.

|  | No additive interaction |  | No multiplicative interaction |  |
| --- | --- | --- | --- | --- |
| Treatment | B = 0 | B = 1 | B = 0 | B = 1 |
| T = 0 | 0.1 | 0.3 | 0.1 | 0.2 |
| T = 1 | 0.5 | 0.7 | 0.3 | 0.6 |

Notional risk of outcome in biomarker negative (B = 0) and biomarker positive (B = 1) patients in the control (T = 0) and experimental treatment group (T = 1) in the scenario of no additive interaction (left) and no multiplicative interaction (right).

Given that interactions on both scales can occur, are relevant and should be analyzed, we need to know how powerful the statistical analyses will be. More specifically, if there is an additive interaction, how likely will this be detected using the “false” model, i.e., the logistic regression? Vice versa, how likely is it to detect a multiplicative interaction when using the linear regression? To answer these questions, we performed a simulation study that will be described in the following.

## Methods

### Simulation framework

In our simulation we start from a population with individuals affected and unaffected by the disease under study, which is indicated by the disease status *D* ∊ {1, 0}. Additional to the general probability of developing the disease, the probability might be influenced by having or having not a certain biomarker status *B* ∊ {1,0}. A random sample *R* of the diseased individuals is recruited to a clinical trial, comparing an experimental treatment with the control treatment, denoted by *T* ∊ {1,0}. The trial aims to answer the research question whether the biomarker *B* is predictive, i.e., whether it modifies the probability of treatment success *y* ∊ {1,0}.

#### Population simulation

We define the prevalence of a dichotomous biomarker *B* by *P*(*B* = 1) = *ϕ*. Populations are simulated by modelling the disease probability by

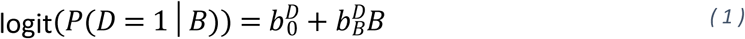

and sampling the disease status *D* from a Bernoulli distribution with probability *P*((*D* = 1|*B*)). Here, 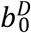 is the baseline log(*odds*) of the disease and 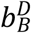 is a prognostic effect of the biomarker *B*.

#### Trial designs

As illustrated in Fig 1, in the “randomize-all” design *n* patients are drawn randomly from a simulated population. Based on the randomization factor *γ* ∊ (0,1), *γn* randomly chosen patients receive the biomarker guided treatment (*T* = 1) and (1 − *γ*)*n* randomly chosen patients receive the control treatment (*T* = 0). After the assignment to a treatment arm the biomarker status is revealed. Thus, the numbers of biomarker positive (*n*_+_) and biomarker negative (*n*_−_) patients in each treatment group are determined by the biomarker prevalence *ϕ*. In the “biomarker-stratified randomization” design the biomarker status is revealed before randomization. This enables to draw *n*_+_ biomarker positive and *n*_−_ biomarker negative, *n* = *n*_+_ + *n*_−_ in total, patients from a simulated population. By specifying *n*_−_ and *n*_+_, the prevalence of the biomarker under consideration is not reflected in this design. In each biomarker stratum, the randomization factors *γ*_+_ ∊ (0,1) and *γ*_−_ ∊ (0,1) determine the proportion of patients receiving control or biomarker guided treatment.

#### Data simulation

In the present simulation study, treatment success is simulated on both the linear and logistic scale in both trial designs for varying parameters. The procedure to simulate this data is as follows:

1. Draw *n* patients from a population based on formula (1).
2. Assign patients to treatment arms based on *γ* or *γ*_+_ and *γ*_−_, depending on the trial design.
3. Calculate the treatment success probability *P*(*y* = 1) by applying eithe

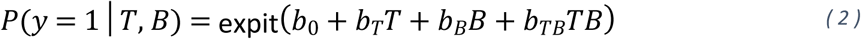 Or

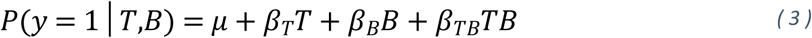

for every patient with 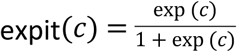, and *T* and *B* denote the treatment and biomarker status, respectively.
4. Sample the treatment success from a Bernoulli distribution using the probability from formula (2) or (3).

We consider *ϕ* ∊ {0.1, 0.25, 0.5} as prevalence for the biomarker, and we use *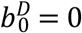* and 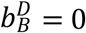 to simulate populations, i.e., there is no prognostic effect of the biomarker. We create study populations of sizes *n* ∊ {100, 200, 500, 1000}. In case of the “biomarker-stratified randomization” trial either half of the study population is biomarker positive and the other half is biomarker negative; alternatively, the proportion of biomarker positive patients is determined by the biomarker prevalence in the respective simulated population, i.e. specifying 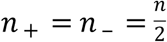 explicitly or specifying only *n*, and from this follows *n*_+_ ≈ *ϕn*. We use *γ*,*γ*_+_,*γ*_−_ ∊ {0.25, 0.5, 0.75} as randomization factors, and in the “biomarker-stratified randomization” trial all combinations of the values of *γ*_+_ and *γ*_−_ are considered. The effect sizes to determine the treatment success probability are the cross-product of a range of possible values. On the linear scale we use

- *β*_0_ = 0.5,
- *β_T_* ∊ {0, 0.1, 0.2, 0.3, 0.4},
- *β_B_* ∊ {−0.4, −0.3, −0.2, − 0.1, 0, 0.1, 0.2, 0.3, 0.4} and
- *β_TB_* ∊ {−0.4, −0.3, −0.2, −0.1, 0, 0.1, 0.2, 0.3, 0.4}.

Combinations of effect sizes leading to a probability of therapy success less than 0 or greater than 1 are excluded, e.g. *β*_0_ = 0.5*, β_T_* = 0, *β_B_* = −0.4*, β_TB_* = −0.4 is not valid.

On the logistic scale we use

- *b*_0_ = 0,
- *b_T_* ∊ {0, 0.2231, 0.4055, 0.5596, 0.6931} corresponding OR ∊ {1,1.25,1.50,1.75,2},
- *b_B_* ∊ { − 0.6931, − 0.5596, − 0.4055, − 0.2231, 0, 0.2231, 0.4055, 0.5596, 0.6931} corresponding to *OR* ∊ {0.5, 0.5713, 0.6667, 0.8, 1, 1.25, 1.5, 1.75, 2}
- *b_TB_* ∊ { − 0.6931, − 0.5596, − 0.4055, − 0.2231, 0, 0.2231, 0.4055, 0.5596, 0.6931} corresponding to *OR* ∊ {0.5, 0.5713, 0.6667, 0.8, 1, 1.25, 1.5, 1.75, 2}.

In total, we use 680 unique effect size combinations for our simulations. Note that effect size combinations having *β_TB_* = 0 or *b_TB_* = 0 act as null models for the respective regression model analysis.

#### Analyses

All simulated data sets are analyzed using both linear and logistic models. Following Kraft et al. [15], the likelihood ratio-based deviance test between the saturated model

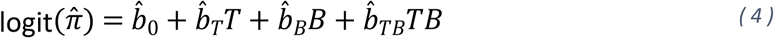

or

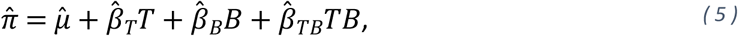

where *π* = *P*(*D* = 1| *B*), and a model considering both main effects of treatment and biomarker but no interaction effect (restricted deviance test) is calculated. In addition, a Wald-like test on the null hypotheses *H*_0_:*b_TB_* = 0 (logistic regression model) or *H*_0_:*β_TB_* = 0 (linear regression model) in the respective saturated models (4) and (5) is performed. To obtain reliable estimates for the power to detect an interaction between treatment and biomarker effect, 1000 replicates are run. For each replicate it is noted whether the two-sided p-value of the respective test is less than *α* = 0.05.

All simulations and analyses are done in R 3.3.1 [16] utilizing the R package batchtools [17]. The code is available in the supplement (S2 Appendix).

## Results

Table 2 shows the estimated frequency of type I errors of the interaction test, i.e., the restricted deviance test, in logistic and linear regression models to detect a interaction effect simulated via the linear (upper part) or logistic (lower part) model. Given are the frequencies in the “randomize-all” trial design with biomarker prevalence *ϕ* = 0.1 and randomization factor *γ* = 0.5 for some selected effect size combinations with no *b_TB_* = log (1) and *β_TB_* = 0 moderate (*b_TB_* = log (1.5) or 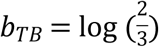 and *β* = ±0.2) and strong (*b_TB_* = log (0.5) or *b_TB_* = log (2) and *β* = ±0.4) effects. The effect sizes are given on both the linear and logistic scale for sample sizes *n*=200 and *n*=500, sorted by the biomarker main effects (Table 2). Other scenarios meeting these restrictions but not displayed are redundant such that the effects *β_T_*,*β_B_*,*b_T_* or *b_B_* have opposite signs or are permuted.

**Table 2.**
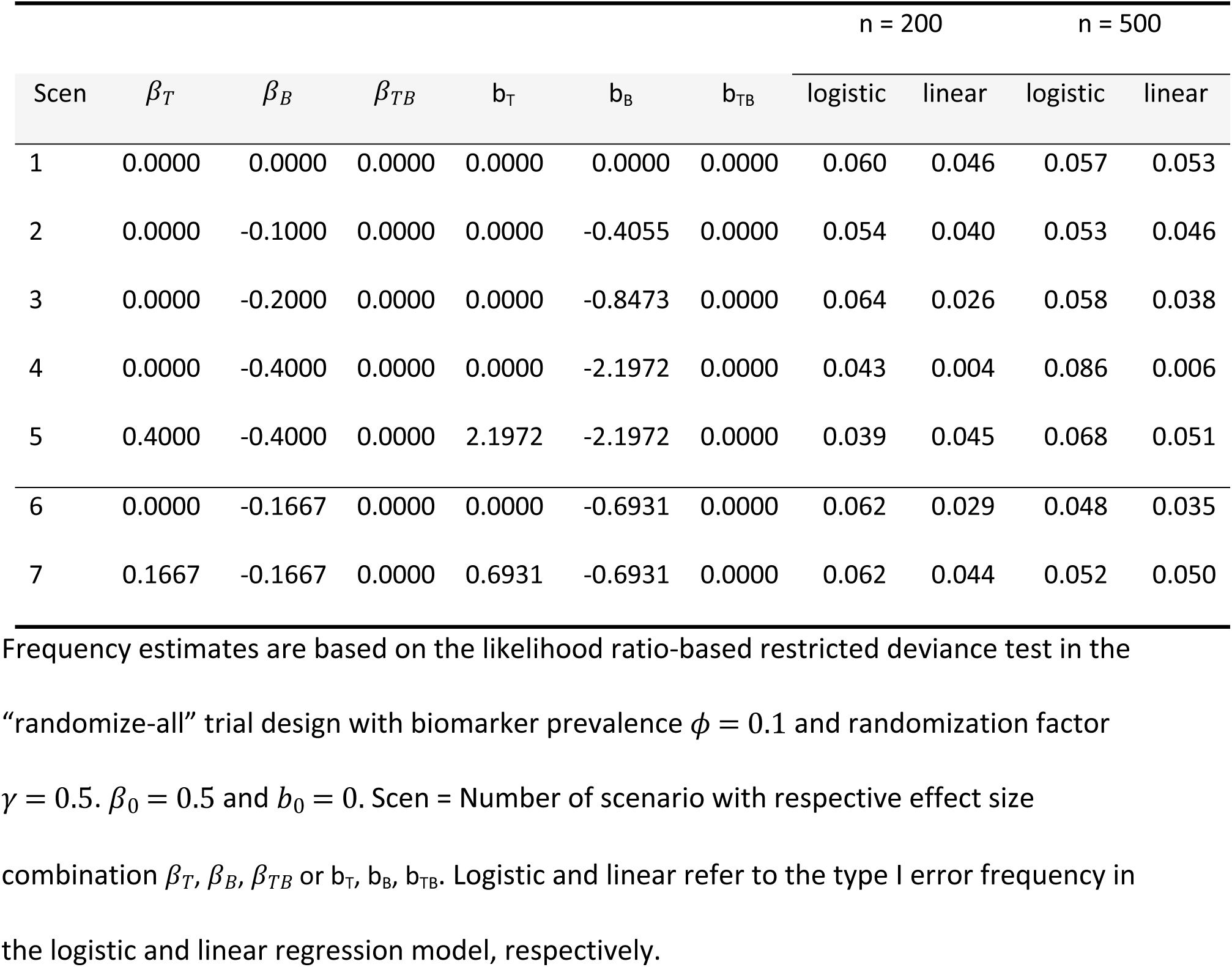
Estimated type I error frequency at the nominal two-sided 0.05 test-level in the “randomize-all” design.

Table 2 shows that the frequency of type I errors for the restricted deviance test in both regression models mainly is near to 0.05, as expected, and thus in line with the specified significance level of α = 0.05. However, in some scenarios the linear and logistic model deviate from the specified significance level. Based on Bradley’s liberal criterion of robustness [18], the type I error frequency should be between 0.025 and 0.075. Both the logistic and the linear model fail to fall into this range in scenario 4, which is characterized by a single strong main effect. The total number and percentage of scenarios violating Bradley’s criterion in the “randomize-all” design is shown in Table 3. In total, 54 times (5% of all scenarios) the logistic model has a type I error outside Bradley’s bounds, whereas the linear model violates this criterion 123 times (11% of all scenarios). Comparing the numbers per model and criterion bound, it is of special interest that the logistic model tends to violate the upper bound (liberal) whereas the linear model tends to violate the lower bound (conservative).

**Table 3.**
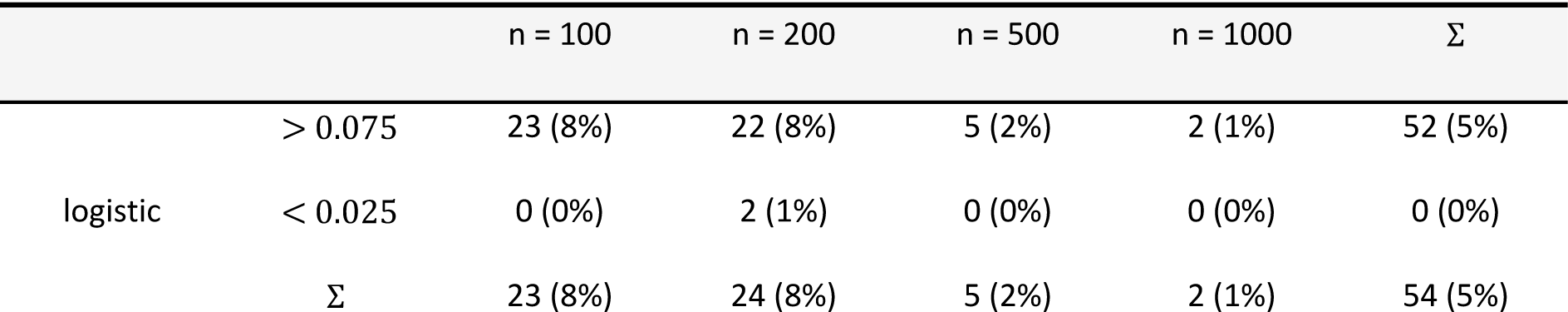
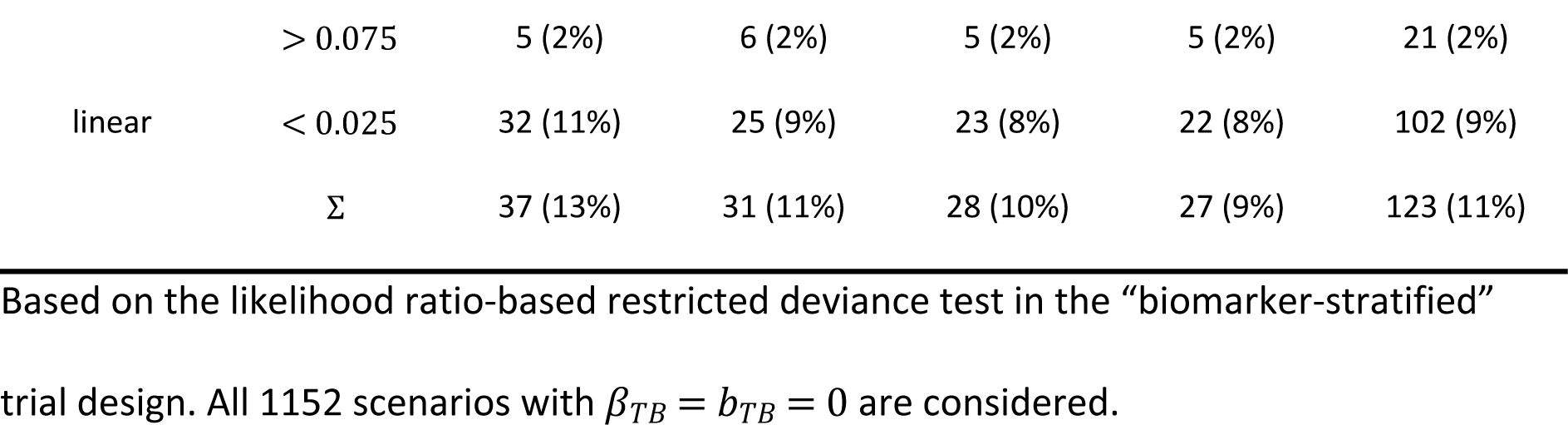
Number of scenarios in which type I error frequencies deviate from Bradley’s criterion [18] in the “randomize-all” design.

|  |  | n = 100 | n = 200 | n = 500 | n = 1000 | Σ |
| --- | --- | --- | --- | --- | --- | --- |
| logistic | > 0.075 | 23 (8%) | 22 (8%) | 5 (2%) | 2 (1%) | 52 (5%) |
|  | < 0.025 | 0 (0%) | 2 (1%) | 0 (0%) | 0 (0%) | 0 (0%) |
|  | Σ | 23 (8%) | 24 (8%) | 5 (2%) | 2 (1%) | 54 (5%) |
| linear | > 0.075 | 5 (2%) | 6 (2%) | 5 (2%) | 5 (2%) | 21 (2%) |
|  | < 0.025 | 32 (11%) | 25 (9%) | 23 (8%) | 22 (8%) | 102 (9%) |
|  | Σ | 37 (13%) | 31 (11%) | 28 (10%) | 27 (9%) | 123 (11%) |

Based on the likelihood ratio-based restricted deviance test in the “biomarker-stratified” trial design. All 1152 scenarios with $\beta_{TB} = b_{TB} = 0$ are considered.

We next look at the power of the restricted deviance test to detect an interaction effect simulated via the linear (Table 4, upper part) or logistic (Table 4, lower part) model in the same setting, i.e., the “randomize-all” trial design with the same effect specifications as before. Results are sorted by the interaction effects.

**Table 4.** Estimated power at the nominal two-sided 0.05 test-level in the “randomize-all” design.

| Scen | $\beta_T$ | $\beta_B$ | $\beta_{TB}$ | $b_T$ | $b_B$ | $b_{TB}$ | n = 200 | | n = 500 | |
| --- | --- | --- | --- | --- | --- | --- | --- | --- | --- | --- |
|  |  |  |  |  |  |  | logistic | linear | logistic | linear |
| 8 | 0.2000 | -0.4000 | 0.0000 | 0.8473 | -2.1972 | 0.5026 | 0.077 | 0.015 | 0.108 | 0.017 |
| 9 | 0.0000 | -0.1000 | 0.1000 | 0.0000 | -0.4055 | 0.4055 | 0.084 | 0.065 | 0.107 | 0.102 |
| 10 | 0.0000 | 0.0000 | -0.1000 | 0.0000 | 0.0000 | -0.4055 | 0.080 | 0.064 | 0.105 | 0.100 |
| 11 | 0.1000 | -0.1000 | -0.1000 | 0.4055 | -0.4055 | -0.4055 | 0.086 | 0.072 | 0.105 | 0.100 |
| 12 | 0.1000 | -0.1667 | -0.1000 | 0.4055 | -0.6931 | -0.4055 | 0.085 | 0.062 | 0.103 | 0.097 |
| 13 | 0.1667 | -0.1667 | -0.1000 | 0.6931 | -0.6931 | -0.4055 | 0.076 | 0.059 | 0.113 | 0.115 |
| 14 | 0.0000 | -0.4000 | 0.2000 | 0.0000 | -2.1972 | 1.3499 | 0.218 | 0.084 | 0.423 | 0.239 |
| 15 | 0.0000 | 0.0000 | -0.2000 | 0.0000 | 0.0000 | -0.8473 | 0.144 | 0.113 | 0.282 | 0.258 |
| 16 | 0.0000 | -0.2000 | -0.2000 | 0.0000 | -0.8473 | -1.3499 | 0.204 | 0.071 | 0.436 | 0.227 |
| 17 | 0.2000 | -0.4000 | -0.2000 | 0.8473 | -2.1972 | -0.8473 | 0.077 | 0.054 | 0.156 | 0.223 |
| 18 | 0.4000 | -0.4000 | -0.2000 | 2.1972 | -2.1972 | -0.8473 | 0.088 | 0.148 | 0.160 | 0.376 |
| 19 | 0.0000 | -0.2000 | 0.4000 | 0.0000 | -0.8473 | 1.6946 | 0.437 | 0.401 | 0.770 | 0.764 |
| 20 | 0.0000 | -0.4000 | 0.4000 | 0.0000 | -2.1972 | 2.1972 | 0.556 | 0.404 | 0.881 | 0.799 |
| 21 | 0.2000 | -0.4000 | 0.4000 | 0.8473 | -2.1972 | 2.1972 | 0.513 | 0.396 | 0.876 | 0.827 |
| 22 | 0.1000 | 0.1000 | -0.0077 | 0.4055 | 0.4055 | 0.0000 | 0.067 | 0.038 | 0.052 | 0.040 |
| 23 | 0.1000 | 0.1667 | -0.0167 | 0.4055 | 0.6931 | 0.0000 | 0.070 | 0.040 | 0.063 | 0.042 |
| 24 | 0.1667 | 0.1667 | -0.0333 | 0.6931 | 0.6931 | 0.0000 | 0.063 | 0.035 | 0.059 | 0.036 |
| 25 | 0.1000 | -0.1667 | -0.0048 | 0.4055 | -0.6931 | 0.0000 | 0.065 | 0.050 | 0.059 | 0.050 |
| 26 | 0.1000 | 0.1000 | 0.0714 | 0.4055 | 0.4055 | 0.4055 | 0.095 | 0.042 | 0.096 | 0.056 |
| 27 | 0.1000 | 0.1667 | 0.0515 | 0.4055 | 0.6931 | 0.4055 | 0.080 | 0.030 | 0.107 | 0.042 |
| 28 | 0.1667 | 0.1667 | 0.0238 | 0.6931 | 0.6931 | 0.4055 | 0.073 | 0.028 | 0.096 | 0.026 |
| 29 | 0.0000 | -0.1667 | 0.0952 | 0.0000 | -0.6931 | 0.4055 | 0.090 | 0.061 | 0.102 | 0.090 |
| 30 | 0.1000 | -0.1667 | 0.0961 | 0.4055 | -0.6931 | 0.4055 | 0.089 | 0.068 | 0.100 | 0.092 |
| 31 | 0.0000 | -0.1000 | -0.0923 | 0.0000 | -0.4055 | -0.4055 | 0.080 | 0.044 | 0.093 | 0.082 |
| 32 | 0.0000 | -0.1667 | -0.0833 | 0.0000 | -0.6931 | -0.4055 | 0.081 | 0.037 | 0.108 | 0.067 |
| 33 | 0.1667 | 0.1667 | -0.1061 | 0.6931 | 0.6931 | -0.4055 | 0.089 | 0.074 | 0.088 | 0.097 |
| 34 | 0.1000 | 0.1000 | 0.1182 | 0.4055 | 0.4055 | 0.6931 | 0.126 | 0.048 | 0.181 | 0.095 |
| 35 | 0.1000 | 0.1667 | 0.0905 | 0.4055 | 0.6931 | 0.6931 | 0.106 | 0.036 | 0.168 | 0.058 |
| 36 | 0.1667 | 0.1667 | 0.0556 | 0.6931 | 0.6931 | 0.6931 | 0.098 | 0.030 | 0.174 | 0.027 |
| 37 | 0.0000 | -0.1000 | 0.1714 | 0.0000 | -0.4055 | 0.6931 | 0.132 | 0.110 | 0.235 | 0.227 |
| 38 | 0.0000 | -0.1667 | 0.1667 | 0.0000 | -0.6931 | 0.6931 | 0.141 | 0.108 | 0.207 | 0.194 |
| 39 | 0.1000 | -0.1667 | 0.1667 | 0.4055 | -0.6931 | 0.6931 | 0.141 | 0.113 | 0.196 | 0.184 |
| 40 | 0.0000 | 0.0000 | -0.1667 | 0.0000 | 0.0000 | -0.6931 | 0.123 | 0.097 | 0.202 | 0.191 |
| 41 | 0.0000 | -0.1000 | -0.1500 | 0.0000 | -0.4055 | -0.6931 | 0.112 | 0.065 | 0.184 | 0.145 |
| 42 | 0.0000 | -0.1667 | -0.1333 | 0.0000 | -0.6931 | -0.6931 | 0.116 | 0.049 | 0.196 | 0.121 |
| 43 | 0.1000 | -0.1000 | -0.1667 | 0.4055 | -0.4055 | -0.6931 | 0.127 | 0.107 | 0.211 | 0.203 |
| 44 | 0.1000 | -0.1667 | -0.1606 | 0.4055 | -0.6931 | -0.6931 | 0.124 | 0.096 | 0.197 | 0.183 |
| 45 | 0.1667 | -0.1667 | -0.1667 | 0.6931 | -0.6931 | -0.6931 | 0.122 | 0.108 | 0.202 | 0.211 |
| 46 | 0.1000 | 0.1000 | -0.1706 | 0.4055 | 0.4055 | -0.6931 | 0.139 | 0.123 | 0.190 | 0.188 |
| 47 | 0.0000 | 0.0000 | -0.4000 | 0.0000 | 0.0000 | -2.1972 | 0.512 | 0.355 | 0.881 | 0.803 |
| 48 | 0.4000 | -0.4000 | -0.4000 | 2.1972 | -2.1972 | -2.1972 | 0.283 | 0.533 | 0.564 | 0.963 |

Power estimates are based on the likelihood ratio-based restricted deviance test in the
“randomize-all” trial design with biomarker prevalence $\phi = 0.1$ and randomization factor
$\gamma = 0.5$ . $\beta_0 = 0.5$ and $b_0 = 0$ . Scen = Number of scenario with respective effect size
combination $\beta_T, \beta_B, \beta_{TB}$ or $b_T, b_B, b_{TB}$ . Logistic and linear refer to the power in the logistic and
linear regression model, respectively.

In some effect size combinations, an interaction effect is present only on one scale. In scenario 8 an interaction effect is present only on the logistic scale. The interaction effect size is rather small compared to the other effect sizes simulated, namely *b_TB_* = 0.5026, rendering an odds ratio of 1.6530. Correspondingly, the power in the logistic regression model to detect the interaction effect is very low at 0.077 (n=200) or 0.108 (n=500). Conversely, scenarios 22 to 25 (Table 4, lower part) reflect the situation of no interaction effect on the logistic scale but only on the linear scale. As in scenario 8 on the logistic scale, the interaction effect sizes are rather small on the linear scale and the power in the linear regression model is very low at 0.035 − 0.05 (n=200) or 0.036 − 0.05 (n=500).

The biggest differences in terms of power between the logistic and linear regression models can be seen if the interaction effect sizes are most extreme and either no or main effects with opposite signs are present. For example, in scenario 48, the restricted deviance test in the linear regression model achieves a power of 0.533, whereas the restricted deviance test in the logistic regression model achieves a power of 0.283 for sample size *n* = 200. This scenario is characterized by a strong negative predictive effect of the biomarker, a positive treatment effect and a strong negative interaction as illustrated in Fig 3A. In other scenarios, the deviance test in the logistic regression model achieves a higher power than in the linear regression model, for example, in scenarios 14, 16, and 20. Here the difference is between ~0.13 and ~0.15, which is illustrated in Fig 3B for scenario 20. These are described by no treatment effects and a negative predictive effect of the biomarker with an additional interaction effect. For all other effect size combinations the differences in terms of power are negligible.

**Fig 3.**
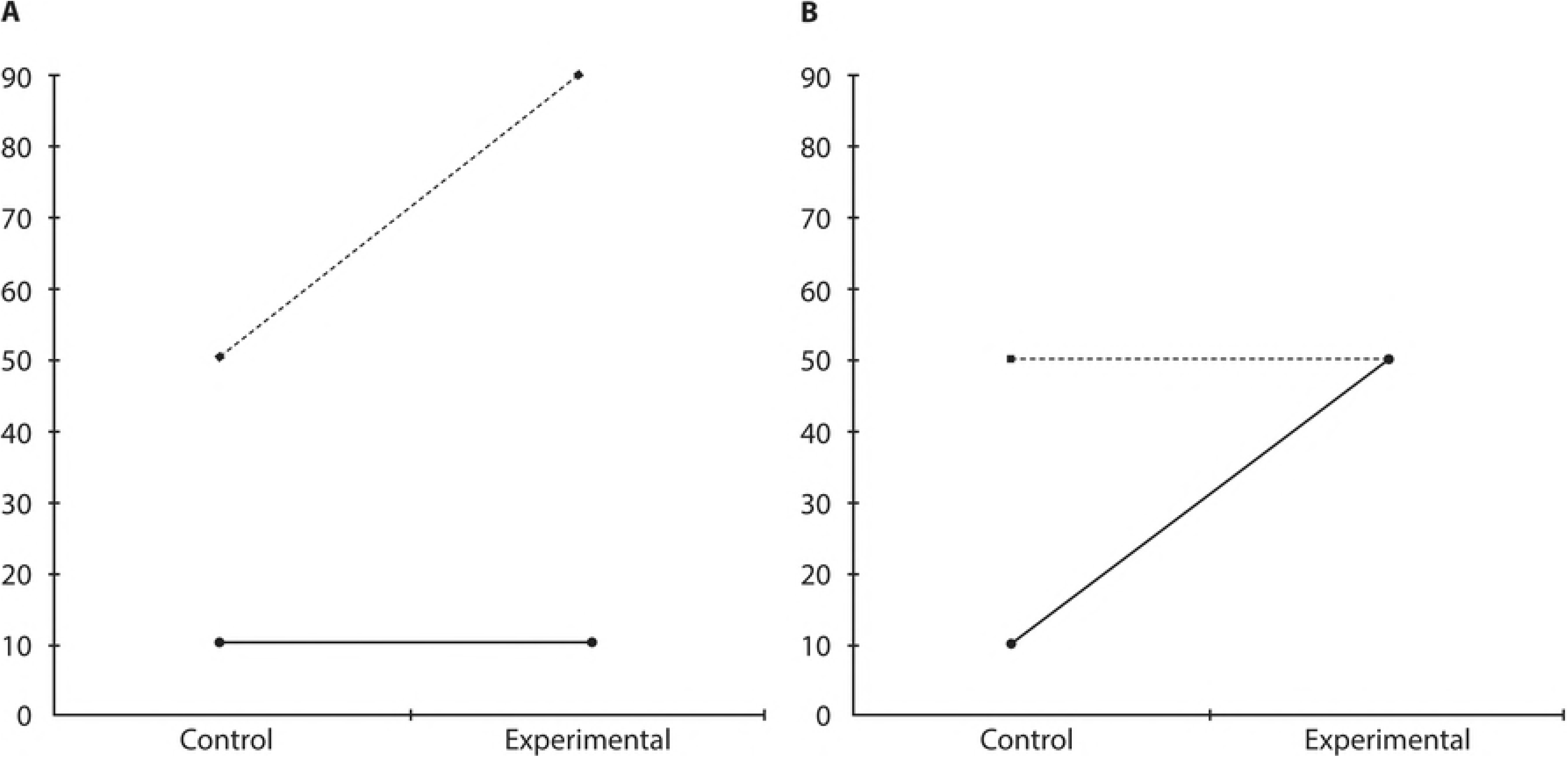
Illustration of scenarios with notable power differences between regression models. Number of patients with a positive outcome. Based on a sample size of 100 in every constellation in (A) scenario 48 characterized by a strong negative predictive effect of the biomarker, a positive treatment effect and a strong negative interaction and in (B) scenario 20 characterized by no treatment effects and a negative predictive effect of the biomarker with an additional interaction effect.

S1 and S2 Tables list the corresponding type I error frequency and estimated power for the same effect size combinations as Tables 2 and 4 in the “biomarker-stratified” trial design with biomarker prevalence *ϕ* = 0.1, randomization factors *γ*_+_ = *γ*_−_ = 0.5, *n*_+_, and *n*_−_ determined by the prevalence of the biomarker *ϕ*. As the same sample sizes are eventually available in the four groups, the estimated frequencies are very similar to those observed in the “randomize-all” trial design. Interestingly, the total number of scenarios violating Bradley’s liberal criterion of robustness in the “biomarker-stratified” design with sample sizes determined by the prevalence of the biomarker (Table 5) is much higher than in the “randomize-all” design (Table 3). Both regression models violate the criterion in about 9% of the scenarios with *β_TB_* = *b_TB_* = 0 (logistic 317 times, linear 309 times). Again, the logistic model tends to be liberal, violating the upper criterion bound, whereas the linear model tends to be conservative, violating the lower criterion bound.

**Table 5.**
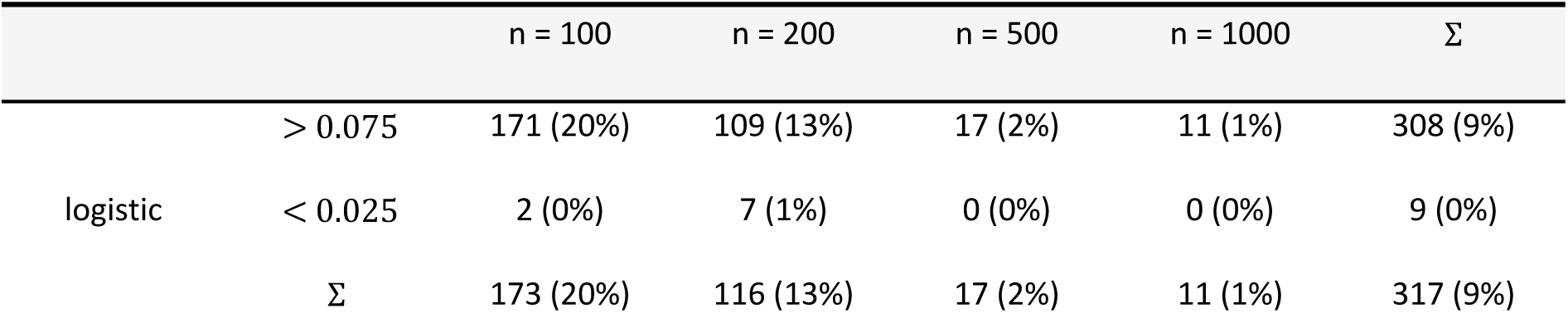
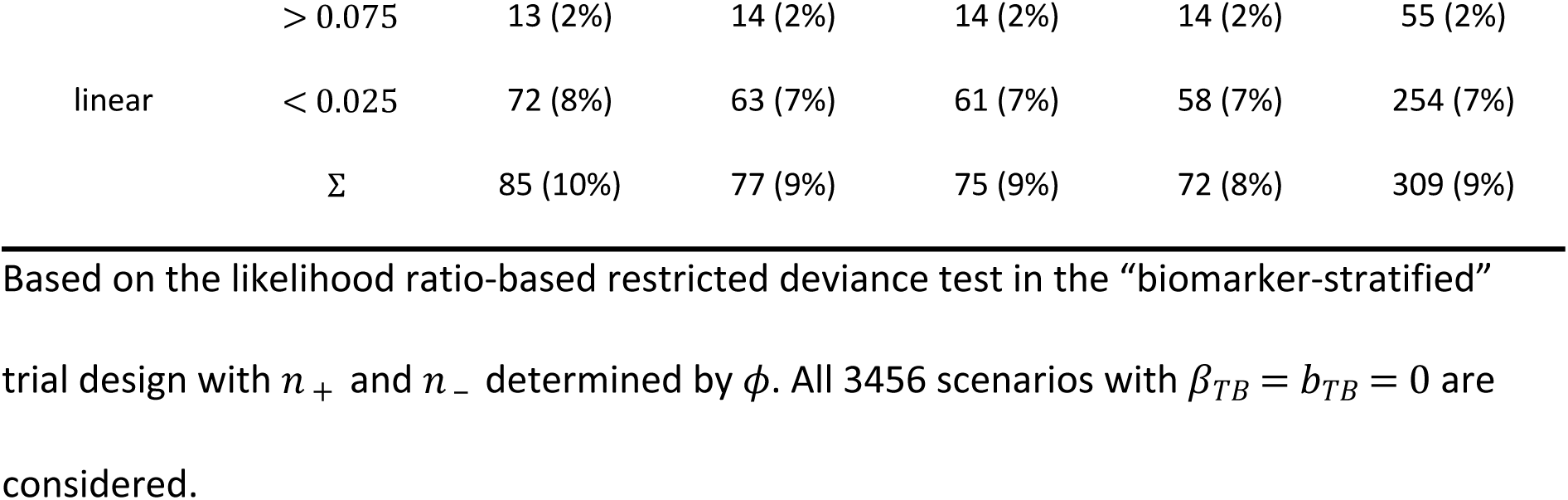
Number of scenarios in which type I error frequencies deviate from Bradley’s criterion [18] in the “biomarker-stratified” design.

Finally, Tables 6, 7 and 8 list the corresponding type I error frequency, scenarios in which the type I error frequencies deviate from Bradley’s criterion, and estimated power for the same effect size combinations with randomization factors *γ*_+_ = *γ*_−_ = 0.5 and fixed proportions of biomarker positive and biomarker negative patients 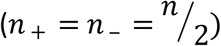. It is therefore assumed that out of a larger patients’ group with biomarker information, only a specified number is selected and included in the trial, so that there is an equal number of biomarker positive and negative cases. In this situation, the estimated type I error is very close to the expected 0.05 in all scenarios with no interaction effect (Table 6), even in scenario 4. Remarkably, in this trial design, the lowest numbers of scenarios violating Bradley’s criterion of robustness is observed (Table 7). The logistic model violates the criterion 36 times and the linear model 81 times, both about 1% of all scenarios with *β_TB_* = *b_TB_* = 0 and *n*_+_, *n*_−_ fixed at 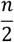. Unexpectedly, in this setting the linear model also tends to be liberal.

**Table 6.**
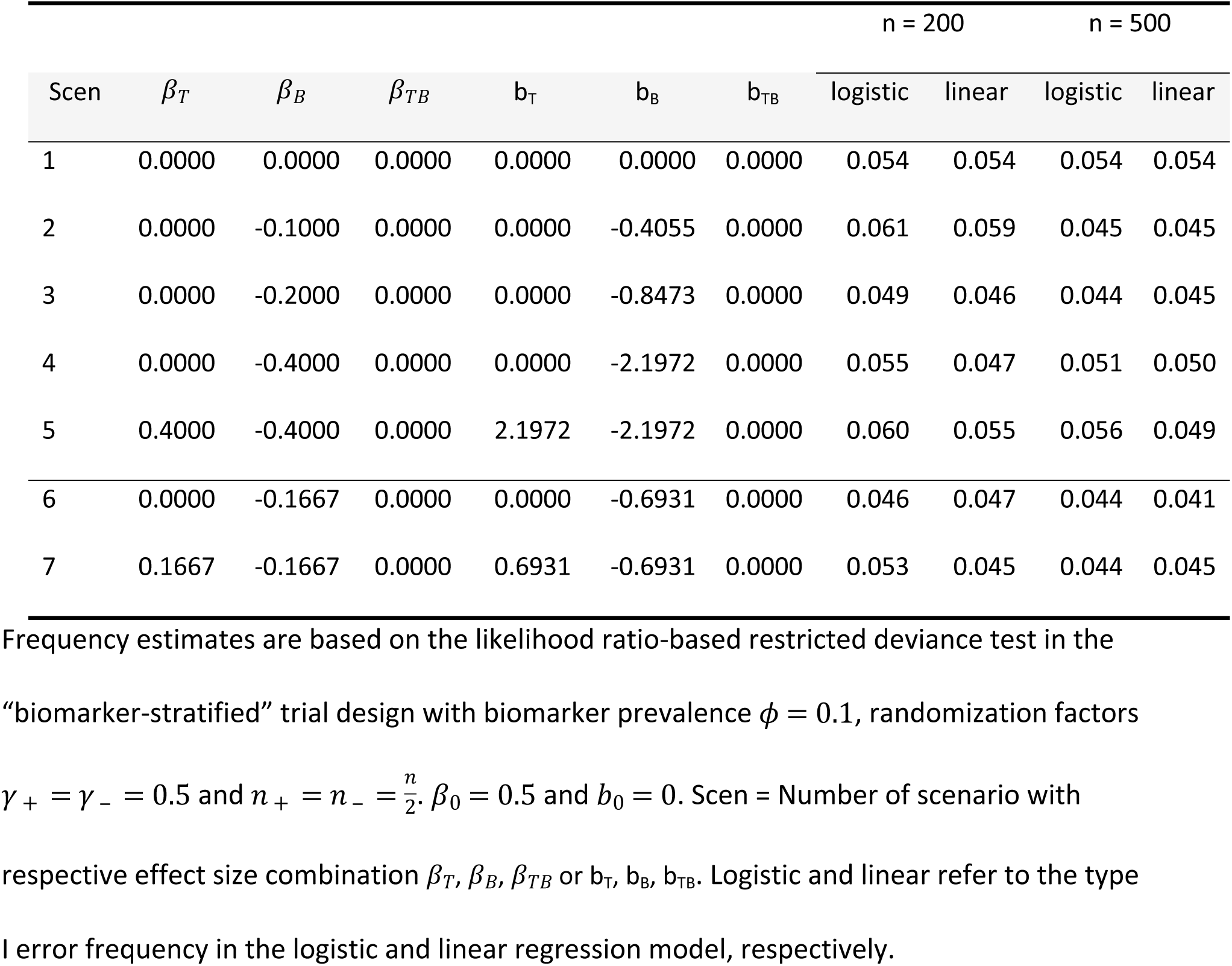
Estimated type I error frequency at the nominal two-sided 0.05 test-level in the “biomarker-stratified” design with fixed proportion of biomarker positive and negative patients.

|  |  |  |  |  |  |  | n = 200 |  | n = 500 |  |
| --- | --- | --- | --- | --- | --- | --- | --- | --- | --- | --- |
| Scen | $\beta_T$ | $\beta_B$ | $\beta_{TB}$ | $b_T$ | $b_B$ | $b_{TB}$ | logistic | linear | logistic | linear |
| 1 | 0.0000 | 0.0000 | 0.0000 | 0.0000 | 0.0000 | 0.0000 | 0.054 | 0.054 | 0.054 | 0.054 |
| 2 | 0.0000 | -0.1000 | 0.0000 | 0.0000 | -0.4055 | 0.0000 | 0.061 | 0.059 | 0.045 | 0.045 |
| 3 | 0.0000 | -0.2000 | 0.0000 | 0.0000 | -0.8473 | 0.0000 | 0.049 | 0.046 | 0.044 | 0.045 |
| 4 | 0.0000 | -0.4000 | 0.0000 | 0.0000 | -2.1972 | 0.0000 | 0.055 | 0.047 | 0.051 | 0.050 |
| 5 | 0.4000 | -0.4000 | 0.0000 | 2.1972 | -2.1972 | 0.0000 | 0.060 | 0.055 | 0.056 | 0.049 |
| 6 | 0.0000 | -0.1667 | 0.0000 | 0.0000 | -0.6931 | 0.0000 | 0.046 | 0.047 | 0.044 | 0.041 |
| 7 | 0.1667 | -0.1667 | 0.0000 | 0.6931 | -0.6931 | 0.0000 | 0.053 | 0.045 | 0.044 | 0.045 |

Frequency estimates are based on the likelihood ratio-based restricted deviance test in the “biomarker-stratified” trial design with biomarker prevalence $\phi = 0.1$ , randomization factors $\gamma_+ = \gamma_- = 0.5$ and $n_+ = n_- = \frac{n}{2}$ , $\beta_0 = 0.5$ and $b_0 = 0$ . Scen = Number of scenario with respective effect size combination $\beta_T, \beta_B, \beta_{TB}$ or $b_T, b_B, b_{TB}$ . Logistic and linear refer to the type I error frequency in the logistic and linear regression model, respectively.

**Table 7.**
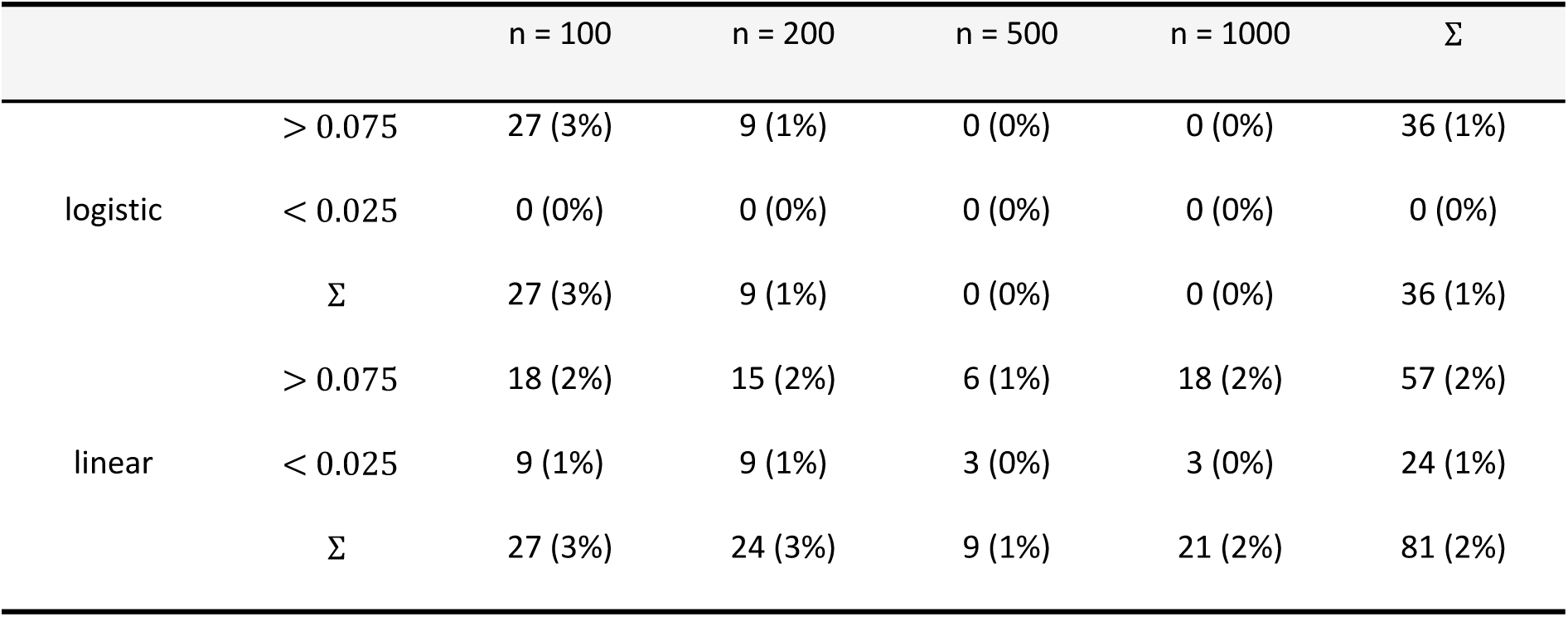
Number of scenarios in which type I error frequencies deviate from Bradley’s criterion [18] in the “biomarker-st ratified” design.

**Table 8.** Estimated power at the nominal two-sided 0.05 test-level in the “biomarker-stratified” design with fixed proportion of biomarker positive and negative patients.

| Scen | $\beta_T$ | $\beta_B$ | $\beta_{TB}$ | $b_T$ | $b_B$ | $b_{TB}$ | n = 200 | | n = 500 | |
| --- | --- | --- | --- | --- | --- | --- | --- | --- | --- | --- |
|  |  |  |  |  |  |  | logistic | linear | logistic | linear |
| 8 | 0.2000 | -0.4000 | 0.0000 | 0.8473 | -2.1972 | 0.5026 | 0.132 | 0.047 | 0.188 | 0.038 |
| 9 | 0.0000 | -0.1000 | 0.1000 | 0.0000 | -0.4055 | 0.4055 | 0.112 | 0.111 | 0.210 | 0.210 |
| 10 | 0.0000 | 0.0000 | -0.1000 | 0.0000 | 0.0000 | -0.4055 | 0.130 | 0.129 | 0.202 | 0.199 |
| 11 | 0.1000 | -0.1000 | -0.1000 | 0.4055 | -0.4055 | -0.4055 | 0.108 | 0.110 | 0.210 | 0.209 |
| 12 | 0.1000 | -0.1667 | -0.1000 | 0.4055 | -0.6931 | -0.4055 | 0.112 | 0.114 | 0.200 | 0.210 |
| 13 | 0.1667 | -0.1667 | -0.1000 | 0.6931 | -0.6931 | -0.4055 | 0.111 | 0.116 | 0.210 | 0.218 |
| 14 | 0.0000 | -0.4000 | 0.2000 | 0.0000 | -2.1972 | 1.3499 | 0.537 | 0.354 | 0.896 | 0.709 |
| 15 | 0.0000 | 0.0000 | -0.2000 | 0.0000 | 0.0000 | -0.8473 | 0.342 | 0.328 | 0.657 | 0.636 |
| 16 | 0.0000 | -0.2000 | -0.2000 | 0.0000 | -0.8473 | -1.3499 | 0.539 | 0.361 | 0.900 | 0.694 |
| 17 | 0.2000 | -0.4000 | -0.2000 | 0.8473 | -2.1972 | -0.8473 | 0.194 | 0.431 | 0.402 | 0.806 |
| 18 | 0.4000 | -0.4000 | -0.2000 | 2.1972 | -2.1972 | -0.8473 | 0.195 | 0.448 | 0.398 | 0.815 |
| 19 | 0.0000 | -0.2000 | 0.4000 | 0.0000 | -0.8473 | 1.6946 | 0.831 | 0.831 | 0.996 | 0.996 |
| 20 | 0.0000 | -0.4000 | 0.4000 | 0.0000 | -2.1972 | 2.1972 | 0.945 | 0.869 | 0.998 | 0.995 |
| 21 | 0.2000 | -0.4000 | 0.4000 | 0.8473 | -2.1972 | 2.1972 | 0.924 | 0.914 | 1.000 | 0.999 |
| 22 | 0.1000 | 0.1000 | -0.0077 | 0.4055 | 0.4055 | 0.0000 | 0.037 | 0.037 | 0.055 | 0.050 |
| 23 | 0.1000 | 0.1667 | -0.0167 | 0.4055 | 0.6931 | 0.0000 | 0.045 | 0.041 | 0.049 | 0.054 |
| 24 | 0.1667 | 0.1667 | -0.0333 | 0.6931 | 0.6931 | 0.0000 | 0.041 | 0.048 | 0.041 | 0.055 |
| 25 | 0.1000 | -0.1667 | -0.0048 | 0.4055 | -0.6931 | 0.0000 | 0.053 | 0.048 | 0.038 | 0.039 |
| 26 | 0.1000 | 0.1000 | 0.0714 | 0.4055 | 0.4055 | 0.4055 | 0.094 | 0.066 | 0.213 | 0.145 |
| 27 | 0.1000 | 0.1667 | 0.0515 | 0.4055 | 0.6931 | 0.4055 | 0.092 | 0.060 | 0.181 | 0.089 |
| 28 | 0.1667 | 0.1667 | 0.0238 | 0.6931 | 0.6931 | 0.4055 | 0.088 | 0.041 | 0.175 | 0.057 |
| 29 | 0.0000 | -0.1667 | 0.0952 | 0.0000 | -0.6931 | 0.4055 | 0.107 | 0.107 | 0.202 | 0.193 |
| 30 | 0.1000 | -0.1667 | 0.0961 | 0.4055 | -0.6931 | 0.4055 | 0.110 | 0.105 | 0.204 | 0.200 |
| 31 | 0.0000 | -0.1000 | -0.0923 | 0.0000 | -0.4055 | -0.4055 | 0.125 | 0.116 | 0.187 | 0.169 |
| 32 | 0.0000 | -0.1667 | -0.0833 | 0.0000 | -0.6931 | -0.4055 | 0.118 | 0.109 | 0.182 | 0.147 |
| 33 | 0.1667 | 0.1667 | -0.1061 | 0.6931 | 0.6931 | -0.4055 | 0.101 | 0.123 | 0.186 | 0.241 |
| 34 | 0.1000 | 0.1000 | 0.1182 | 0.4055 | 0.4055 | 0.6931 | 0.205 | 0.135 | 0.442 | 0.305 |
| 35 | 0.1000 | 0.1667 | 0.0905 | 0.4055 | 0.6931 | 0.6931 | 0.179 | 0.103 | 0.401 | 0.195 |
| 36 | 0.1667 | 0.1667 | 0.0556 | 0.6931 | 0.6931 | 0.6931 | 0.154 | 0.057 | 0.366 | 0.107 |
| 37 | 0.0000 | -0.1000 | 0.1714 | 0.0000 | -0.4055 | 0.6931 | 0.235 | 0.236 | 0.520 | 0.520 |
| 38 | 0.0000 | -0.1667 | 0.1667 | 0.0000 | -0.6931 | 0.6931 | 0.235 | 0.228 | 0.491 | 0.482 |
| 39 | 0.1000 | -0.1667 | 0.1667 | 0.4055 | -0.6931 | 0.6931 | 0.216 | 0.212 | 0.505 | 0.505 |
| 40 | 0.0000 | 0.0000 | -0.1667 | 0.0000 | 0.0000 | -0.6931 | 0.249 | 0.244 | 0.498 | 0.483 |
| 41 | 0.0000 | -0.1000 | -0.1500 | 0.0000 | -0.4055 | -0.6931 | 0.235 | 0.213 | 0.476 | 0.419 |
| 42 | 0.0000 | -0.1667 | -0.1333 | 0.0000 | -0.6931 | -0.6931 | 0.212 | 0.179 | 0.427 | 0.339 |
| 43 | 0.1000 | -0.1000 | -0.1667 | 0.4055 | -0.4055 | -0.6931 | 0.231 | 0.227 | 0.474 | 0.475 |
| 44 | 0.1000 | -0.1667 | -0.1606 | 0.4055 | -0.6931 | -0.6931 | 0.216 | 0.216 | 0.467 | 0.473 |
| 45 | 0.1667 | -0.1667 | -0.1667 | 0.6931 | -0.6931 | -0.6931 | 0.228 | 0.237 | 0.451 | 0.482 |
| 46 | 0.1000 | 0.1000 | -0.1706 | 0.4055 | 0.4055 | -0.6931 | 0.249 | 0.247 | 0.488 | 0.488 |
| 47 | 0.0000 | 0.0000 | -0.4000 | 0.0000 | 0.0000 | -2.1972 | 0.939 | 0.871 | 1.000 | 0.998 |
| 48 | 0.4000 | -0.4000 | -0.4000 | 2.1972 | -2.1972 | -2.1972 | 0.718 | 0.972 | 0.979 | 1.000 |

Power estimates are based on the likelihood ratio-based restricted deviance test in the “biomarker-stratified” trial design with biomarker prevalence $\phi = 0.1$ , randomization factors $\gamma_+ = \gamma_- = 0.5$ and $n_+ = n_- = \frac{n}{2}$ , $\beta_0 = 0.5$ and $b_0 = 0$ . Scen = Number of scenario with respective effect size combination $\beta_T, \beta_B, \beta_{TB}$ or $b_T, b_B, b_{TB}$ . Logistic and linear refer to the power in the logistic and linear regression model, respectively.

Based on the likelihood ratio-based restricted deviance test in the “biomarker-stratified” trial design with 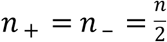. All 3456 scenarios with *β_TB_* = *b_TB_* = 0 are considered.

Similar as in the previous designs, if an interaction effect is present only on one scale, it is hard to detect, resulting in a low power. In general, however, the pattern of the estimated power is very similar to before, with an overall higher power due to balanced sample sizes.

For an overview, Table 9 shows a comparison of the estimated power across the considered scenarios. Here, the number of scenarios is given in which the power in the linear and logistic regression model is comparable (less than 3% difference), in which one of the models is slightly better (difference between 3% and 10%), and in which one of the models is better (difference greater than 10%). These numbers are given for all considered scenarios and only for scenarios without extreme effect constellations. For the vast majority of scenarios, the difference in estimated power of the linear and logistic model is irrelevant, i.e., the difference is less than 3%, and differences are smaller with larger sample sizes. If relevant power differences are observed, this is usually in favor of the logistic model. Interestingly, this pattern remains the same when scenarios with extreme effect combinations are not considered.

**Table 9.** Power comparison for restricted deviance test.

|  |  | Randomize-All |  | Biomarker-Stratified |  | Biomarker-Stratified* |  |
| --- | --- | --- | --- | --- | --- | --- | --- |
|  |  | n=200 | n=500 | n=200 | n=500 | n=200 | n=500 |
| all scenarios | logistic >> linear | 24 | 6 | 23 | 4 | 2 | 2 |
| (599) |  | (4.0%) | (1.0%) | (3.8%) | (0.7%) | (0.3%) | (0.3%) |
|  | logistic > linear | 232 | 78 | 184 | 77 | 34 | 13 |
|  |  | (38.7%) | (13.0%) | (30.7%) | (12.9%) | (5.7%) | (2.2%) |
|  | logistic = linear | 332 | 499 | 379 | 503 | 550 | 576 |
|  |  | (55.4%) | (83.3%) | (63.3%) | (84.0%) | (91.8%) | (96.2%) |
|  | logistic < linear | 11 | 16 | 13 | 15 | 12 | 8 |
|  |  | (1.8%) | (2.7%) | (2.2%) | (2.5%) | (2.0%) | (1.3%) |
|  | logistic << linear | 0 | 0 | 0 | 0 | 1 | 0 |
|  |  | (0%) | (0%) | (0%) | (0%) | (0.2%) | (0%) |
| excluding most | logistic >> linear | 24 | 6 | 23 | 4 | 2 | 2 |
| extreme scenarios |  | (4.5%) | (1.1%) | (4.3%) | (0.7%) | (0.4%) | (0.4%) |
| (535) | logistic > linear | 212 | 75 | 164 | 75 | 32 | 13 |
|  |  | (39.3%) | (13.9%) | (30.4%) | (13.9%) | (5.9%) | (2.4%) |
|  | logistic = linear | 297 | 450 | 343 | 453 | 498 | 516 |
|  |  | (55.1%) | (83.5%) | (63.6%) | (84.0%) | (92.4%) | (95.7%) |
|  | logistic < linear | 6 | 8 | 9 | 7 | 6 | 8 |
|  |  | (1.1%) | (1.5%) | (1.7%) | (1.3%) | (1.1%) | (1.5%) |
|  | logistic << linear | 0 | 0 | 0 | 0 | 1 | 0 |
|  |  | (0%) | (0%) | (0%) | (0%) | (0.2%) | (0%) |
| excluding extreme | logistic >> linear | 24 | 6 | 23 | 4 | 2 | 2 |
| scenarios |  | (4.7%) | (1.2%) | (4.5%) | (0.8%) | (0.4%) | (0.4%) |
| (515) | logistic > linear | 206 | 74 | 157 | 74 | 32 | 13 |
|  |  | (40.0%) | (14.4%) | (30.5%) | (14.4%) | (6.2%) | (2.5%) |
|  | logistic = linear | 282 | 428 | 331 | 431 | 474 | 492 |
|  |  | (54.8%) | (83.1%) | (64.3%) | (83.7%) | (92.0%) | (95.5%) |
|  | logistic < linear | 3 | 7 | 4 | 6 | 6 | 8 |
|  |  | (0.6%) | (1.4%) | (0.8%) | (1.2%) | (1.2%) | (1.6%) |
|  | logistic << linear | 0 | 0 | 0 | 0 | 1 | 0 |
|  |  | (0%) | (0%) | (0%) | (0%) | (0.2%) | (0%) |

The above results were obtained from using the likelihood-based restricted deviance test for interaction. Using a Wald-like test instead produces the same results in the linear model, but lower type I and type II errors in the logistic model. The number of scenarios in which the type I error frequencies deviate from Bradley’s criterion in the Wald-like test are shown in S3 to S5 Tables. In addition, we presented only a limited selection of the simulation results, but the preceding descriptions are also valid for the other simulation settings, and a compilation of all results can be found in S6 Table (note that the numbers of the effect size combinations in S6 Table are not the same as in Tables 2, 4, 6, 8).

## Discussion and conclusions

The predictiveness of a biomarker can be evaluated via the treatment-by-biomarker interaction in linear or logistic regression models for a binary outcome, and we have derived the relationship between the effects of the linear model and the logistic model (S1 Appendix). The translation between ORs from the logistic and AARs from the linear model might be useful, since the ARRs can in turn be used to calculate the NNT which is helpful for the clinical interpretation. In a comprehensive simulation study, we compared the power of the linear and logistic regression models to detect the predictiveness of a biomarker under a variety of scenarios in the randomize-all and the biomarker-stratified design. In general, we found that the differences in power to detect interaction were minor. Visible differences in power were detected in rather unrealistic scenarios of effect size combinations and were usually in favor of the logistic model. If the number of biomarker-positive and biomarker-negative patients in the biomarker-stratified design was guided by the prevalence of the biomarker, we did not find notable differences compared to the randomize-all design. However, if equal subgroups of biomarker-positive and biomarker-negative patients could be selected in the biomarker-stratified design, the power was decidedly greater owing to the balanced samples sizes.

Different baseline probabilities were not considered in our simulations. These could have impact on the power of both regression models and the power differences as well, especially if they are close to 0 and 1. However, we assume that these values only play a minor role in applications.

For choosing between the logistic and the linear model for a clinical trial that aims at showing predictiveness of a biomarker one should therefore consider the following factors:

1. The linear regression model has statistical disadvantages. For example, the predicted probability might be out of the 0-1-range of possible values. Furthermore, the model fit is rather poor if the predicted probabilities are close to 0 or 1. In the logistic regression model, the error terms follow a binomial distribution, and statistical properties are generally good for a binary outcome [19].
2. As expected, the type I error frequency was adequate in both models, unless the scenarios were extreme, where the linear model was sometimes conservative.
3. Power was comparable, again unless the effect size combinations were highly unusual. If there were differences, the logistic model usually had higher power than the linear probability model.
4. The effects from the linear model can be interpreted in a more straightforward way, which was also pointed out be Hellevik [14] in the case of main effects, and ARR and OR can be translated into each other.

Thus, the choice of the appropriate regression model should always be driven by the primary aim of a study [19] and is influenced by two different currents, the statistical properties and the ease of interpretation. From the statistical viewpoint one should favor the most sparse model. Following this, one could estimate both models and select the one with the least number of non-zero estimates. However, our simulations have shown that it is hard to find effect size combinations with non-zero effects on only one scale. Thus, from a practical point of view one should favor the logistic regression model, and inference based on the logistic regression model estimates should be theoretically more valid than inference based on linear regression model estimates. Consequently, the logistic model should be used if the presence of an interaction effect is to be tested. Concerning the interpretation regarding the treatment effect in different groups, the linear model seems recommendable. With our results in mind, it therefore seems recommendable to estimate logistic regression models because of their statistical properties, test for interaction effects and calculate and report both ARRs and ORs from these using the formulae provided in the appendix.

## Supporting information

**S1 Appendix. Relation between absolute risk reductions from linear probability models and odds ratios from logistic regression models.**

**S2 Appendix. Simulation code.** Refer to included README for further information.

**S1 Table. Estimated type I error frequency at the nominal two-sided 0.05 test-level in the “biomarker-stratified” design with biomarker prevalence 0.1.** Frequency estimates are based on the likelihood ratio-based restricted deviance test in the “biomarker-stratified” trial design with biomarker prevalence *ϕ* = 0.1, randomization factors *γ*_+_ = *γ*_−_ = 0.5 and *n*_+_ and *n*_−_ are determined by *ϕ*. *β*_0_ = 0.5 and *b*_0_ = 0. Scen = Number of scenario with respective effect size combination *β_T_*, *β_B_*, *β_TB_* or b_T_, b_B_, b_TB_. Logistic and linear refer to the type I error frequency in the logistic and linear regression model, respectively.

**S2 Table. Estimated power at the nominal two-sided 0.05 test-level in the “biomarker-stratified” design with biomarker prevalence 0.1.** Power estimates are based on the likelihood ratio-based restricted deviance test in the “biomarker-stratified” trial design with biomarker prevalence *ϕ*=0.1, randomization factors *γ*_+_ = *γ*_−_ = 0.5 and *n*_+_ and *n*_−_ are determined by *ϕ*.*β*_0_=0.5 and *b*_0_=0. Scen = Number of scenario with respective effect size combination *β_T,_ β_B,_ β_TB_* or b_T_, b_B_, b_TB_. Logistic and linear refer to the power in the logistic and linear regression model, respectively.

**S3 Table. Number of scenarios in which type I error frequencies deviate from Bradley’s criterion [18] in the “randomize-all” design.** Based on the Wald-test in the “biomarker-stratified” trial design. All 1152 scenarios with are *β_TB_* = *b_TB_* = 0 considered.

**S4 Table. Number of scenarios in which type I error frequencies deviate from Bradley’s criterion [18] in the “biomarker-stratified” design.** Based on the Wald-test in the “biomarker-stratified” trial design with *n*_+_ and *n*_−_ determined by *ϕ*. All 3456 scenarios with *β_TB_* = *b_TB_* =0 are considered.

**S5 Table. Number of scenarios in which type I error frequencies deviate from Bradley’s criterion [18] in the “biomarker-stratified” design.** Based on the Wald-test in the “biomarker-stratified” trial design with 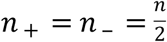. All 3456 scenarios with *β_TB_* = *b_TB_* = 0 are considered.

**S6 Table. Compilation of all simulation results.** The numbers of the effect size combinations are not the same as in Tables 2, 4, 6, 8.

